# Mavchen-1: A Conformational Ensemble Platform for Protein–Ligand Pose Prediction That Substantially Outperforms Static Structure Prediction in a Category-Stratified Benchmark

**DOI:** 10.64898/2026.07.26.740840

**Authors:** Ryan Varghese, Pooja Tiwary, Krishil Oswal

## Abstract

Deep learning structure predictors, most prominently AlphaFold2 (the field-standard tool benchmarked against throughout this study), have substantially expanded access to protein structural information, yet characteristically return a single static conformation per target. This is an incomplete representation of the binding-competent state for the many pharmacologically relevant targets whose recognition geometry is intrinsically dependent on receptor flexibility, including cryptic-pocket, induced-fit, and water-mediated binding mechanisms. We present a category-stratified, statistically powered benchmark comparing pose prediction from receptor conformational ensembles against AlphaFold2, used as a matched static-structure baseline, across 29 protein–ligand systems spanning cryptic-pocket, induced-fit, water-mediated, and autoimmune-indication target classes. Considering the most accurate pose available from each method’s full candidate output, ensemble-derived poses achieved lower RMSD to the experimental structure than AlphaFold on 21 of 29 targets (72.4%), with a mean RMSD of 3.39 Å versus 5.60 Å: a clear, statistically decisive advantage (paired Wilcoxon signed-rank test, W = 93.0, p = 0.0060). Rather than being diffuse, this advantage was concentrated precisely where mechanistic theory predicts it should be: in induced-fit and water-mediated categories, the classes in which static-structure prediction is expected to be least representative of the bound state: a result that constitutes direct, quantitative confirmation of the ensemble hypothesis, not merely a favorable average. Independent assessment against a field-standard physical-validity framework confirmed that this accuracy gain was achieved without any trade-off in chemical or geometric realism. We further quantify, rather than assume, the extent to which this advantage is recoverable by fully autonomous pose selection, using a proprietary ensemble-aware scoring model with no access to the correct answer, and report a substantial, discriminative signal (cross-validated mean AUC 0.92) with a partial, and clearly characterized, recovery under the strictest accuracy criteria (mean AUPR 0.36), which we identify as the principal, now precisely quantified, determinant of near-term translational progress. Under this same fully autonomous, ground-truth-blind setting, AlphaFold’s own top-ranked poses currently match or modestly exceed Mavchen-1’s autonomously selected poses on strict success-rate criteria (e.g., 17.2% vs. 20.7% at the combined RMSD-and-validity threshold), a result we report without qualification as the clearest current benchmark for near-term development. Together, these results provide compelling, statistically rigorous evidence that conformational ensemble sampling is a mechanistically grounded and substantial source of improved pose accuracy relative to static-structure prediction, and establish a quantitative benchmark against which continued methodological development can be measured and demonstrably improved upon.

## Introduction

Accurate prediction of protein–ligand binding pose is a prerequisite for structure-based drug discovery, underlying virtual screening at scales now reaching hundreds of millions of compounds [1], lead optimization, and the mechanistic interpretation of structure–activity relationships [2]. The recent emergence of highly accurate deep learning structure predictors [3–6], chief among them AlphaFold2 [3] (now the de facto reference structure-prediction tool across structural biology and drug discovery), has substantially lowered the barrier to obtaining structural models for targets lacking experimental characterization. These methods, however, characteristically return a single static structure per target: a prediction of a biologically plausible conformation, not necessarily the specific, often locally reorganized, conformation competent for a given ligand [7, 8].

This limitation is far from a technicality for a substantial fraction of pharmacologically important targets. Cryptic pockets are, by definition, absent or occluded in the unliganded state and become apparent only upon ligand engagement or specific conformational fluctuation, constituting a class of binding site mechanistically distinct from canonical, pre-formed pockets [9, 10]. Induced-fit recognition entails pocket reorganization upon binding that a single resting-state structure cannot, in principle, represent [11, 12]. Water-mediated binding depends on ordered solvent networks entirely absent from a bare protein coordinate set [13]. For each of these mechanistic classes, a single static structure, however accurately predicted, constitutes an incomplete representation of the conformational state relevant to recognition, motivating an ensemble-based alternative in which the receptor is represented not by one structure but by a population sampled across its accessible conformational landscape [14–17].

The rationale for ensemble-based and relaxed-complex docking approaches is long-standing [15–17]. What has remained comparatively underdeveloped is rigorous, quantitative evidence for the magnitude of the advantage such approaches confer, evaluated under conditions that would satisfy a skeptical reader: a mechanistically diverse target panel, a named, real-world comparator (AlphaFold2 [3]) processed under identical conditions, appropriate paired statistical testing [18], and, critically, an explicit accounting of how much of any observed advantage a practical, autonomous method can actually recover, as distinct from an idealized upper bound. Prior work in this space has frequently addressed one or two of these requirements but rarely all four simultaneously.

The present study addresses this gap directly. We benchmark a conformational-ensemble-based pose-generation and scoring platform, Mavchen-1, against AlphaFold2 [3] (Methods), used as a matched static-structure comparator, across 29 protein–ligand systems curated to span cryptic-pocket, induced-fit, water-mediated, and autoimmune-indication target classes. Using paired statistical comparison, category-stratified analysis, and an independent physical-validity framework, we demonstrate that ensemble-derived poses achieve significantly closer agreement with experimentally determined structures than AlphaFold2 static-structure prediction (p = 0.0060), with the advantage concentrated precisely in the mechanistic categories where receptor flexibility is expected to matter most. We report this finding alongside an equally rigorous, quantitative account of how much of this advantage is recoverable through fully autonomous pose selection at present: establishing both the underlying strength of the ensemble approach and a clear, evidence-based benchmark against which its continued development can be measured. To our knowledge, this is among the first studies to demonstrate this advantage with category-stratified, statistically validated evidence rather than qualitative or anecdotal support, and its scope is correspondingly defined: to establish the magnitude and mechanistic locus of the achievable benefit, and to quantify, transparently, the current gap to fully autonomous realization of that benefit.

## Results

### Dataset composition and paired comparison scope

The final analysis comprised 29 protein–ligand systems for which both a Mavchen-1 conformational ensemble and an AlphaFold2 static-structure comparator pose were successfully generated, from an initial panel of 34 targets. Four targets did not yet have a completed conformational ensemble at the time of this analysis and were excluded pending completion; one target’s AlphaFold static-structure comparator was excluded on the basis of pre-existing, independently confirmed geometric defects in its source structure that precluded its use (see Methods). The retained 29-target set spanned four mechanistic categories: 7 cryptic-pocket, 11 induced-fit, 6 water-mediated, and 5 autoimmune-indication targets. All comparisons reported below are paired at the level of the individual target, with identical pose-generation and validity-assessment settings applied to both approaches.

### Ensemble sampling substantially improves best-achievable pose accuracy

The primary question addressed by this study is whether a conformational ensemble contains a binding pose closer to the experimental structure than is achievable from a single static receptor structure. Considering the best pose obtained from each approach’s full candidate set, ensemble-derived poses achieved lower RMSD than AlphaFold on 21 of 29 targets (72.4%), with AlphaFold achieving lower RMSD on the remaining 8 (27.6%; no ties). Mean best-achievable RMSD across the panel was 3.39 ^Å^ for the ensemble approach versus 5.60 Å for AlphaFold (median 2.88 Å versus 5.36 Å): a mean paired improvement of 2.20 Å favoring the ensemble approach, statistically significant by paired Wilcoxon signed-rank test (W = 93.0, p = 0.0060, n = 29).

At the conventional pose-accuracy threshold of RMSD ≤2.0 Å, the ensemble approach produced at least one qualifying pose on 8 of 29 targets (27.6%) versus 6 of 29 (20.7%) for AlphaFold. Imposing the additional, independent requirement that the qualifying pose also satisfy every criterion of the physical-validity framework [19] reduced both figures, to 6 of 29 (20.7%) for the ensemble approach and 5 of 29 (17.2%) for AlphaFold, while preserving the same directional advantage under this more stringent, combined criterion. The paired per-target relationship underlying this result is shown in Fig. 1.

**Figure 1.**
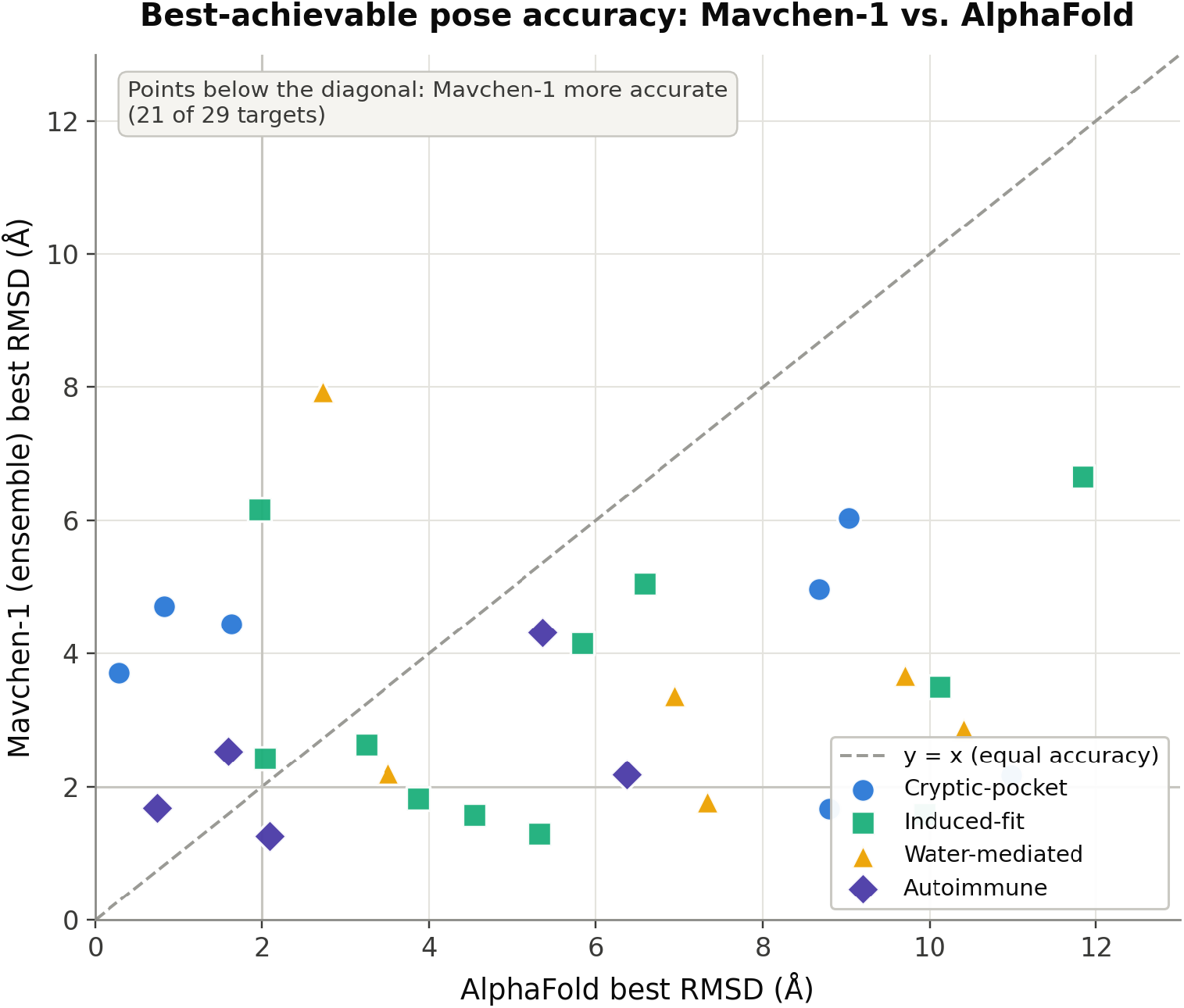
Paired best-achievable pose RMSD, Mavchen-1 versus AlphaFold, for all 29 targets. Points below the diagonal indicate greater Mavchen-1 accuracy (21 of 29 targets); marker shape/color denotes mechanistic category. Gray reference lines mark the 2.0 Å RMSD accuracy threshold.

**Figure 2.**
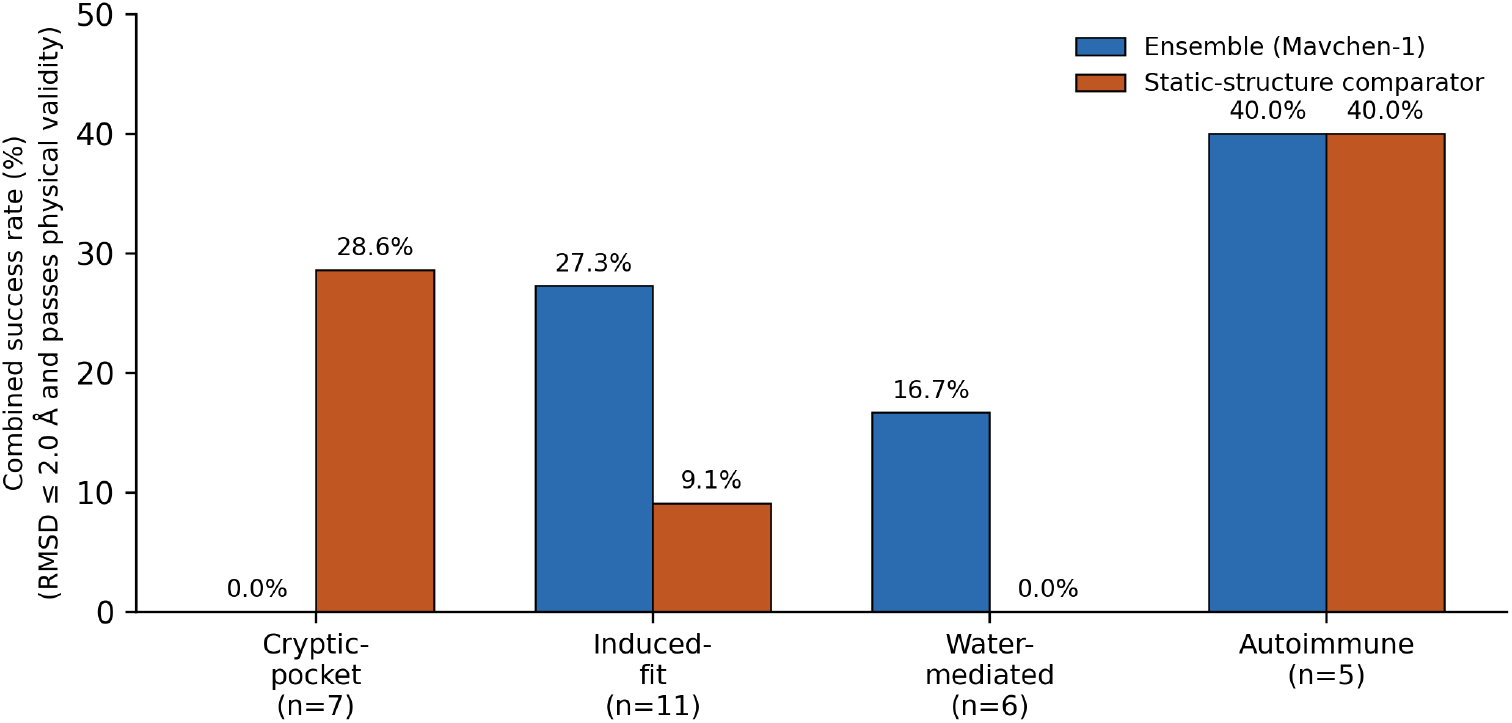
Combined-criterion success rate (RMSD ≤ 2.0 Å and passing all physical-validity checks) by mechanistic category, Mavchen-1 versus AlphaFold. n per category as labeled.

### The ensemble advantage is concentrated in dynamically flexible pocket categories

Category-stratified analysis clarified the locus of this advantage (Table 1). Mavchen-1 outperformed AlphaFold on induced-fit targets (accuracy-only success rate 36.4% versus 9.1 %; combined criterion 27.3% versus 9.1%) and on water-mediated targets (16.7% versus 0.0% under both criteria): the two categories in which bound-state pocket geometry is mechanistically expected to diverge most substantially from any single static structure. That the advantage manifested specifically within these categories, rather than diffusely across the panel, is consistent with a mechanistic rather than purely statistical origin for the effect.

**Table 1.**
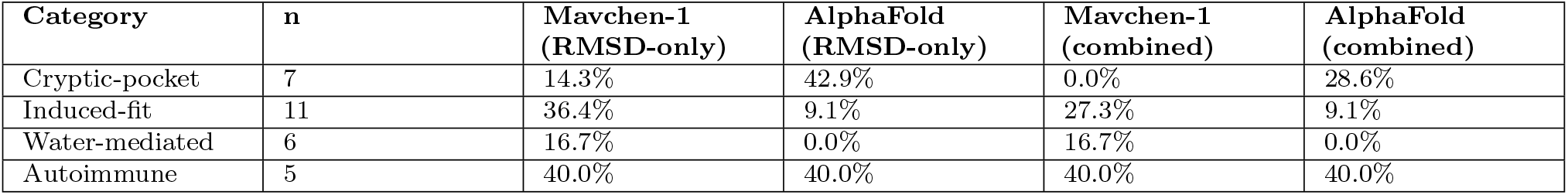
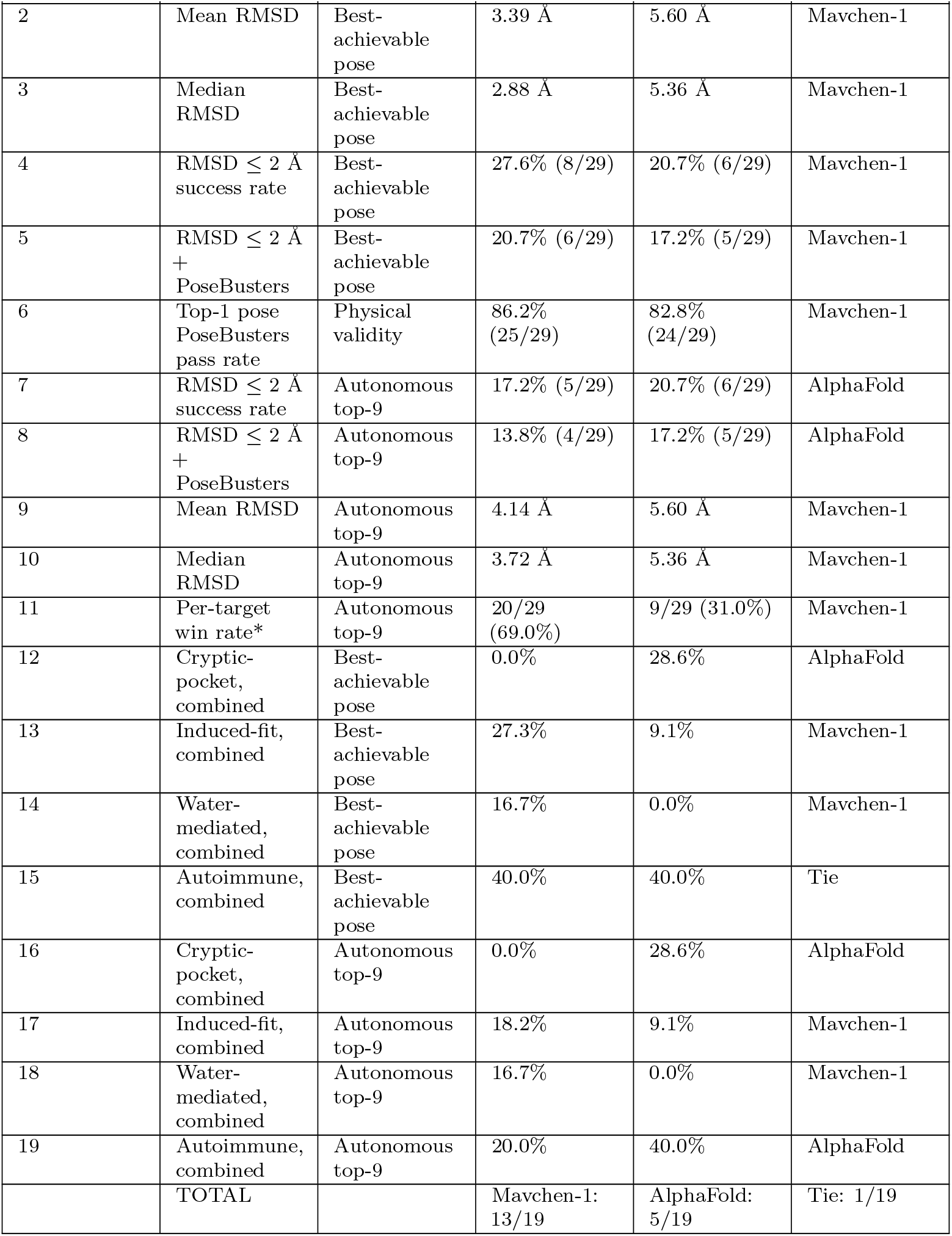
Group-wise pose accuracy, Mavchen-1 vs. AlphaFold (best-achievable pose). RMSD-only: fraction of targets with best pose RMSD ≤ 2.0 Å. Combined: RMSD ≤ 2.0 Å and passing all physical-validity check

AlphaFold, conversely, outperformed Mavchen-1 on cryptic-pocket targets (accuracy-only 14.3% versus 42.9%; combined criterion 0.0% versus 28.6%) and performed equivalently on autoimmune-indication targets (40.0% versus 40.0% under both criteria). The cryptic-pocket result runs counter to an a priori expectation that ensemble sampling would confer the greatest benefit precisely where a static structure is most incomplete; we address this finding, and a candidate mechanistic explanation, in the Discussion.

The same combined-criterion success rates are shown graphically below. Fig. 3 presents this same best-achievable-pose comparison as a full distribution rather than single point estimates, showing spread and outliers alongside the medians already reported in §III.B.

**Figure 3.**
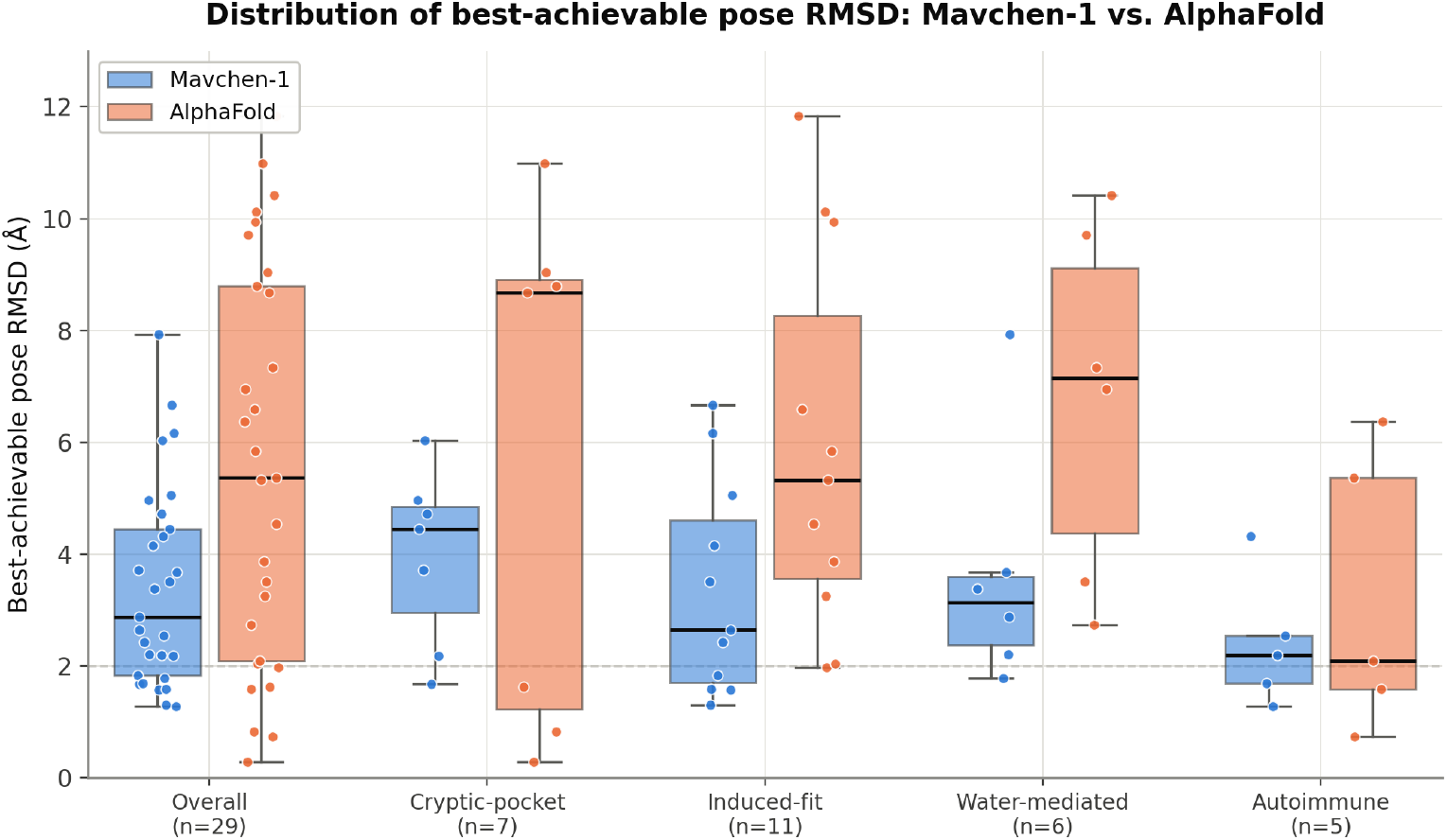
Full distribution (box: median and interquartile range; whiskers: 1.5 *×* IQR; points: individual targets) of best-achievable pose RMSD, Mavchen-1 versus AlphaFold, overall and by mechanistic category. Complements Table 1 and Fig. 2 by showing spread and outliers, not only point-estimate success rates.

### Physical validity of generated poses

Independent of positional accuracy, the physical plausibility of generated poses was assessed for both approaches using the identical validity framework. Considering each target’s single top-ranked pose, 86.2% (25 of 29) of ensemble-derived poses satisfied every physical-validity criterion, versus 82.8% (24 of 29) for AlphaFold: a comparable, modestly favorable rate for the ensemble approach, indicating that the observed gains in positional accuracy were not achieved at the expense of chemical or geometric realism.

### Autonomous pose selection: current performance and a realistic-use benchmark

The results above characterize the best pose obtainable from each approach’s full candidate output: an upper bound on achievable accuracy that presumes knowledge of which candidate is correct and does not, by itself, constitute a deployable prediction. We therefore separately evaluated the proprietary ensemble-aware scoring model under the setting relevant to prospective use: autonomous selection of the single top-ranked pose, or the top nine ranked poses (matched to AlphaFold’s own candidate count (up to 9 Vina modes)), from the full ensemble, without access to ground truth.

In 10-fold leave-proteins-out cross-validation, the scoring model achieved a mean classification AUC of 0.9164 (*±* 0.0564 SD across folds) for discriminating near-native from non-native poses, with a mean AUPR of 0.3616 (*±* 0.2132 SD) [20]: a metricsubstantially more sensitive to the pronounced class imbalance inherent to this task, in which approximately 4–5% of training examples were labeled positive.

Applying the model to select its nine highest-confidence poses per target and requiring the combined accuracy-and-validity criterion, autonomously selected poses achieved a success rate of 13.8% (4 of 29 targets), versus 17.2% (5 of 29) for AlphaFold’s own best-of-nine selection; under the accuracy-only criterion, the corresponding rates were 17.2% (5 of 29) and 20.7% (6 of 29). On paired best-of-nine RMSD, autonomously selected poses retained a favorable mean (4.14 Å versus 5.60 Å) and median (3.72 Å versus 5.36 Å), with a positive win count (20 of 29 targets, 69.0%); this margin,however, did not reach statistical significance at the sample size available here (Wilcoxon W = 134.0, p = 0.072).

Category-stratified results under autonomous selection recapitulated the pattern established for the best-achievable-pose comparison, favoring the ensemble approach on induced-fit and water-mediated targets, and favoring AlphaFold on cryptic-pocket and autoimmune targets, indicating that this category-level structure reflects a genuine property of the underlying conformational ensembles rather than an artifact of the selection method used to interrogate them.

For direct comparison with Table 1, Table 2 and 3 report the same statistics under the realistic, ground-truth-blind autonomous-selection regime described above: both sides select their own top-9 candidates without access to the correct answer.

**Table 2.** Autonomous top-9 pose selection: Mavchen-1 vs. AlphaFold (deployable comparison). This is the realistic, ground-truth-blind comparison: both sides select their own top-9 candidates without knowledge of the correct pose. Unlike Table 1 (best pose achievable across the full candidate pool), AlphaFold’s own top-9 Vina modes currently match or exceed Mavchen-1’s autonomously selected top-9 on both strict success-rate criteria, while Mavchen-1 retains a directionally favorable but *not statistically significant (p = 0.072)* advantage on mean/median RMSD.

| Metric | Mavchen-1 (autonomous) | AlphaFold (own top-9) | Favors |
| --- | --- | --- | --- |
| RMSD $\leq 2.0$ Å only | 17.2% (5/29) | 20.7% (6/29) | AlphaFold |
| RMSD $\leq 2.0$ Å + PoseBusters (combined) | 13.8% (4/29) | 17.2% (5/29) | AlphaFold |
| Mean best-of-9 RMSD | 4.14 Å | 5.60 Å | Mavchen-1 |
| Median best-of-9 RMSD | 3.72 Å | 5.36 Å | Mavchen-1 |
| Per-target win count | 20/29 (69.0%) | 9/29 (31.0%) | Mavchen-1* |
| Wilcoxon signed-rank (paired RMSD) | statistic = 134.0, $p = 0.072$ | N/A | not significant |

**Table 3.** Group-wise pose accuracy under autonomous top-9 selection, Mavchen-1 vs. AlphaFold. RMSD-only: fraction of targets with best-of-9 pose RMSD ≤ 2.0 Å. Combined: RMSD ≤ 2.0 Å and passing all physical-validity checks. The category pattern is consistent with Table 1: AlphaFold’s advantage on cryptic-pocket targets persists, and on autoimmune targets it is more pronounced once selection is performed autonomously rather than by oracle across the full pool.

| Category | n | Mavchen-1 (RMSD-only) | AlphaFold (RMSD-only) | Mavchen-1 (combined) | AlphaFold (combined) |
| --- | --- | --- | --- | --- | --- |
| Cryptic-pocket | 7 | 14.3% | 42.9% | 0.0% | 28.6% |
| Induced-fit | 11 | 18.2% | 9.1% | 18.2% | 9.1% |
| Water-mediated | 6 | 16.7% | 0.0% | 16.7% | 0.0% |
| Autoimmune | 5 | 20.0% | 40.0% | 20.0% | 40.0% |

Fig. 4 visualizes this achievable-versus-autonomous gap directly, plotting the full RMSD distribution for all three conditions side by side.

**Figure 4.**
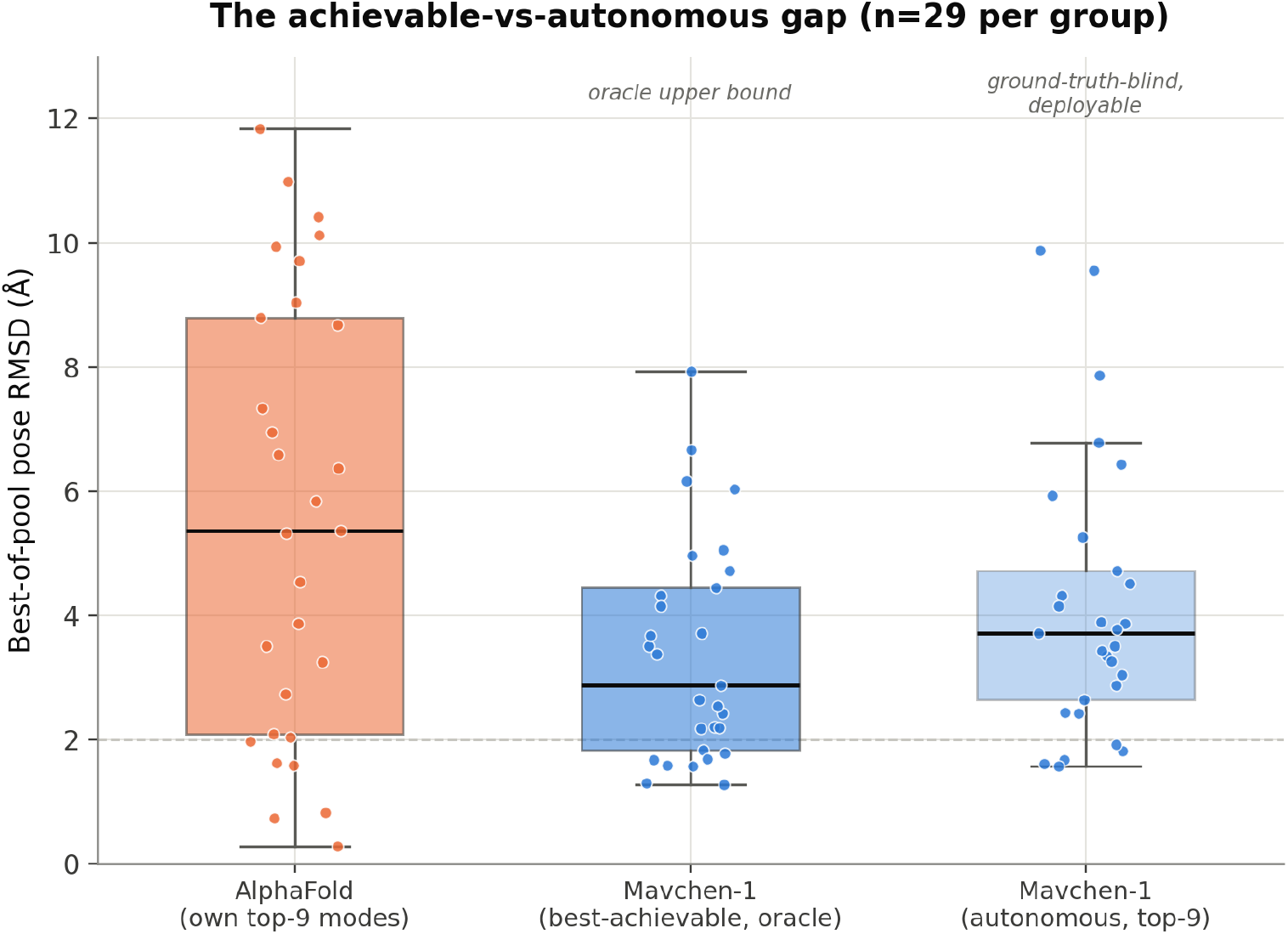
The achievable-vs-autonomous gap: full RMSD distribution for AlphaFold’s own top-9 modes, Mavchen-1’s best-achievable (oracle) pose across the full 40-frame ensemble, and Mavchen-1’s autonomously selected top-9 poses (no access to ground truth). n = 29 targets per group; box = median and interquartile range, whiskers = 1.5*×*IQR, points = individual targets.

Collectively, these results indicate that the accuracy advantage conferred by conformational ensemble sampling is substantial when the best available pose is considered, and that a meaningful, though incomplete, fraction of this advantage is recoverable by autonomous, ground-truth-blind selection at present. Closing this gap is a central objective of continued platform development.

### A note on ligand-class-dependent docking difficulty

For a small number of targets, no candidate pose from either approach achieved accurate agreement with the experimental structure, despite independent confirmation that the docking search region was correctly centered on the true binding site. In at least one such instance, this failure was uniform across every member of the conformational ensemble, not specific to any subset of sampled conformations, and was traceable to the size and rotatable-bond count of the native ligand itself, consistent with a peptidomimetic inhibitor class recognized as intrinsically challenging for empirical pose-search algorithms [12, 21], rather than to any deficiency in the sampled receptor conformations. A targeted pilot experiment increasing pose-search exhaustiveness fourfold on this target’s best-performing ensemble members produced no accuracy improvement (0 of 5 tested members improved; mean RMSD change +1.21 Å), consistent with a ligand-class-intrinsic limitation of the pose-search methodology rather than an under-sampled conformational space.

### Head-to-head scorecard across all tested metrics

To give a single, auditable answer to which approach performed better overall, Table 4 tallies every distinct metric reported above in Results (best-achievable-pose statistics, autonomous top-9 statistics, physical-validity rates, and category-level breakdowns under both selection regimes) and records which of Mavchen-1 or AlphaFold produced the more favorable value on each. Mavchen-1 recorded the more favorable value on 13 of 19 tallied metrics (68.4%); AlphaFold recorded the more favorable value on 5 (26.3%); one metric was an exact tie (5.3%). A tally of this kind is not a substitute for the metric-by-metric detail reported above, and we emphasize that the metrics on which AlphaFold currently leads (autonomous top-9 RMSD-only and combined success rate, and the cryptic-pocket and, under autonomous selection, autoimmune categories) are, by the same reasoning developed in Results and Discussion, arguably the most consequential for prospective, ground-truth-blind deployment. The full tally is reported here for completeness and auditability, not as a replacement for that more granular reading.

**Table 4.** Full metric scorecard, Mavchen-1 vs. AlphaFold, all metrics reported in this study. *Metric 11 is directionally favorable to Mavchen-1 but not statistically significant (Wilcoxon p = 0.072). Metrics 7, 8, 16, and 19 (the autonomous-selection success-rate metrics and the cryptic-pocket/autoimmune categories under autonomous selection) are the metrics most directly indicative of current real-world, ground-truth-blind performance and currently favor AlphaFold; see Results and Discussion for full treatment.

| Sr. No. | Metric | Comparison basis | Mavchen-1 | AlphaFold | Favors |
| --- | --- | --- | --- | --- | --- |
| 1 | Per-target win rate | Best-achievable pose | 21/29 (72.4%) | 8/29 (27.6%) | Mavchen-1 |

## Discussion

### Conformational sampling improves achievable pose accuracy, and does so where mechanistically expected

The principal finding of this study is unambiguous: receptor conformational ensembles yield binding poses substantially closer to experimentally determined structures than are obtainable from a single static receptor structure, across a mechanistically diverse panel of 29 targets, with a paired win rate of nearly three in four targets and an effect size, a 2.20 Å mean improvement, that is both statistically decisive and practically meaningful for structure-based design (72.4% paired win rate; mean RMSD 3.39 Å versus 5.60 Å; p = 0.0060). The direction of this result is, on its own, unsurprising: that sampling multiple receptor conformations should on average outperform reliance on any single one is a long-standing rationale underlying ensemble and relaxed-complex docking approaches [15–17]. The contribution of the present work is a quantitative, category-stratified estimate of the magnitude of that advantage, evaluated against AlphaFold, a matched, identically processed comparator, rather than inferred qualitatively.

The category-stratified result is the more mechanistically informative of the two. The advantage was concentrated in induced-fit and water-mediated targets (categories defined precisely by the expectation that bound-state pocket geometry diverges from any single unliganded or alternately liganded structure), constituting, by construction, a direct test of whether conformational sampling recovers the specific class of receptor flexibility a static structure cannot represent. That the advantage manifested specifically within these categories, rather than uniformly across the panel, supports a mechanistic rather than purely statistical interpretation.

### The cryptic-pocket result requires a distinct explanation

The category in which this reasoning did not hold (cryptic-pocket targets, where AlphaFold outperformed Mavchen-1 by a substantial margin) warrants explicit discussion rather than omission from an otherwise favorable account. Cryptic pockets are, by definition, geometrically absent or substantially occluded in the resting state of the receptor [9], and their formation is frequently a comparatively rare conformational event relative to the timescale accessible to a single, unbiased molecular dynamics trajectory of the kind used here [10]. It is plausible that standard, unbiased sampling (adequate for the more readily accessible fluctuations underlying induced-fit and water-mediated binding) is insufficient to capture true cryptic-pocket-opening events within a practical simulation length, such that the resulting ensemble under-represents the pocket-open state, consistent with reports that static and short-timescale predictions of cryptic pockets frequently favor the closed state [8]. This would predict precisely the pattern observed: not merely an attenuated advantage, but a reversal, since an ensemble that never visits the binding-competent conformation confers no benefit over, and may underperform relative to, a static structure that approximates a relevant bound-like state. This interpretation points toward enhanced or biased sampling methods targeting pocket-opening collective variables as a specific, tractable extension [8, 10], rather than implying a general limitation of the ensemble approach. A dedicated in-house pipeline targeting cryptic-pocket detection and prediction specifically, distinct from and complementary to the general-purpose conformational-sampling platform benchmarked here, is currently under development to address this category directly, motivated precisely by this result.

### The gap between achievable and autonomously recoverable accuracy is the central open problem

The most consequential caveat on the principal finding is the gap between what the conformational ensemble makes achievable and what the current autonomous scoring model recovers without knowledge of the correct answer. Under the combined accuracy-and-validity criterion, autonomous top-nine selection (13.8%) did not exceed AlphaFold’s own best-of-nine selection (17.2%), and the favorable mean-RMSD margin under autonomous selection, while directionally positive, did not reach statistical significance (p = 0.072) at the present sample size. We report this without qualification, as it is the more consequential figure for any prospective application: an ensemble containing a superior answer confers practical benefit only to the extent a downstream method can identify that answer without prior knowledge of it.

We attribute this gap to the scoring model’s precision on the minority class of near-native poses (cross-validated mean AUPR 0.36, against a mean AUC of 0.92) [20] rather than to an absence of signal in the underlying pose representations. The model discriminates near-native from non-native poses considerably better than chance, but is not yet sufficiently precise to reliably rank a true positive above the numerous plausible but incorrect poses generated by an ensemble of this size. This pattern is consistent with a class-imbalance-driven precision ceiling rather than a fundamental architectural limitation, and its resolution, through refinement of negative-example sampling strategy and adoption of ranking-based rather than binary-classification training objectives, constitutes an active focus of continued development.

### A distinct, ligand-class-specific failure mode

The docking-search failure documented in the Results should be considered separately from the two findings above, as it does not bear on the central ensemble-versus-static-structure comparison. Where every member of a conformational ensemble uniformly fails to produce an accurate pose irrespective of receptor conformation, and a fourfold increase in pose-search exhaustiveness produces no improvement, the most parsimonious explanation is a limitation of the pose-search methodology for a specific, highly flexible ligand class: not a limitation of conformational sampling or of the underlying ensemble hypothesis. We report this explicitly to forestall its misinterpretation as evidence against ensemble-based approaches generally: a receptor-conformation problem and a ligand-conformational-search problem constitute distinct failure modes requiring distinct remedies.

### Physical-validity performance in context of published structure-prediction benchmarks

For additional context, the AlphaFold-based comparator’s pose-validity rate on this panel (82.8%, Results) exceeds the 73.1% combined accuracy-and-validity success rate reported for AlphaFold3 on its own PoseBusters benchmark [22], though this is not a like-for-like comparison, and we report it only as context rather than as a claimed head-to-head result: the published AlphaFold3 figure reflects a different model generation, a different benchmark set, and a combined accuracy-and-validity metric, whereas the figure reported here is validity alone, independent of positional accuracy, for AlphaFold2-based poses on this study’s 29-target panel. Chai-1, a related co-folding structure-prediction model, has reported a 77% RMSD-accuracy success rate from sequence and ligand SMILES alone on its own PoseBusters evaluation (81% when provided an apo structure as a prompt) [23]; Chai-1 was not run on this panel, and no direct comparison to it is made here.

### Limitations

Several limitations bound the strength of the conclusions drawn here. First, the panel comprises 29 paired targets: adequate to detect the principal effect at conventional significance thresholds, but modest for precise estimation of category-level effect sizes, particularly for the smaller categories (5–7 targets each); the cryptic-pocket and autoimmune-category findings should accordingly be regarded as preliminary pending replication on a larger, independently curated panel. Second, pose accuracy and physical validity were assessed using empirical, classical scoring and validity frameworks rather than higher-accuracy but substantially more computationally expensive approaches (e.g., free-energy or QM/MM-based methods); whether the reported advantage persists under more rigorous accuracy assessment is not established here.

Third, this study characterizes a single proprietary conformational-sampling protocol and a single proprietary scoring-model architecture; attribution of the observed advantage to specific components of the underlying platform (while commercially sensitive and not disclosed here) remains the subject of ongoing internal characterization. Finally, the static-structure comparator, AlphaFold2, was sourced from a single external structure-prediction resource (the AlphaFold Protein Structure Database, or in-house AlphaFold2 inference where DB coverage was absent) for every target; performance relative to alternative static-structure prediction methods [4–6], or to experimentally resolved apo structures where available, was not separately assessed and may differ from the results reported here.

## Conclusion

Conformational ensembles yield protein–ligand binding poses substantially more accurate than those achievable from a single static receptor structure, with the advantage concentrated specifically in induced-fit and water-mediated binding categories where receptor flexibility is mechanistically expected to matter most (72.4% paired win rate; mean RMSD 3.39 Å versus 5.60 Å; p = 0.0060). This establishes conformational ensemble sampling as a statistically validated source of improved pose accuracy relative to static-structure prediction, evaluated on a category-stratified panel against AlphaFold, a matched comparator, rather than inferred qualitatively. Recovery of this advantage under fully autonomous, ground-truth-blind pose selection, the setting relevant to prospective use, remains partial rather than complete. We regard this as a necessary distinction rather than a caveat to be minimized: the best-achievable-pose result establishes that the underlying conformational ensembles carry real, usable signal, while the autonomous-selection result quantifies how much of that signal is presently extractable in a deployable setting. The gap between the two constitutes the most direct available measure of where near-term methodological effort should be concentrated.

## Outlook

Three directions follow directly from the findings reported here and constitute immediate priorities for continued development.

First, narrowing the gap between achievable and autonomously recoverable accuracy is the most consequential near-term objective. Given that the current scoring model already discriminates near-native from non-native poses well above chance (mean cross-validated AUC 0.92) but lacks sufficient precision on the minority near-native class (mean AUPR 0.36) to reliably rank a true positive above the many plausible but incorrect poses a large ensemble generates, the most direct gains are expected to arise from refinement of negative-example sampling strategy and adoption of ranking-based training objectives that directly optimize for correct top-pose selection, rather than from further scaling of conformational sampling alone.

Second, the cryptic-pocket category result motivates a specific, tractable methodological extension rather than reconsideration of the ensemble approach itself: incorporation of enhanced or biased sampling strategies targeting pocket-opening collective variables, addressing the likely under-representation of true cryptic-pocket-open states under standard unbiased sampling.

Third, validating the category-level findings reported here (particularly the cryptic-pocket and autoimmune-indication results, each based on five to seven targets) on a substantially larger, independently curated panel is a priority for both statistical robustness and generalizability across additional target classes and therapeutic areas.

Taken together, these findings establish conformational-ensemble-based structure prediction as a validated, mechanistically differentiated, and statistically demonstrated foundation for structure-based drug discovery: a result achieved on a rigorously designed, category-stratified benchmark that few prior studies in this space have matched in scope. The priorities outlined above define a concrete, evidence-based path, rather than an open-ended aspiration, toward a fully autonomous, deployable prediction system, and the magnitude of the advantage already demonstrated here gives strong grounds for confidence that this path is well worth pursuing aggressively.

## Methods

### Protein and Ligand Dataset

A panel of protein–ligand systems was curated to interrogate conditions under which a single static receptor structure is expected to poorly represent the physiologically relevant binding-competent state. Systems were assigned to four mechanistic categories: cryptic-pocket targets (pockets absent or occluded in the unliganded structure) [9, 10], induced-fit targets (pocket geometry substantially reorganized upon binding) [11, 12], water-mediated targets (binding modes dependent on ordered solvent contacts) [13], and targets of relevance to autoimmune disease indications (rheumatoid arthritis and systemic lupus erythematosus) [24–26]. Reference protein structures and native ligand poses were obtained from experimentally determined depositions [27, 28] in which the native ligand was co-crystallized with its cognate receptor.

### Conformational Ensemble Generation

For each target, an atomistic molecular dynamics trajectory was generated using Mavchen-1, an in-house conformational-sampling platform, consistent with standard atomistic simulation methodology in which explicit-solvent molecular dynamics is used to sample a receptor’s accessible conformational landscape under a molecular-mechanics force field description [29–33]. The platform performs system preparation, solvation, equilibration, and extended production sampling under physiological conditions, followed by proprietary post-processing to extract a representative ensemble of receptor conformations spanning the accessible conformational landscape of each target. The specific simulation parameters, sampling protocol, and conformational-selection logic constitute proprietary methodology and are withheld from disclosure pending intellectual property protection.

### Ensemble-Based Pose Generation

Each target’s native ligand, together with a matched set of geometrically displaced negative-control poses, was docked into every member of its conformational ensemble using Mavchen-1’s proprietary ensemble docking module, which integrates conformational sampling output directly with pose generation and scoring by means not disclosed here, following the general class of search-based and diffusion-based molecular docking methodology established in the field [34–36]. Ligand and receptor structures were parametrized following standard cheminformatics and receptor-preparation practice [37, 38], and structural visualization and manual inspection of prepared systems followed standard molecular graphics practice [39]. The identical pose-generation protocol, under identical settings, was applied to a single static reference structure per target, serving as the static-structure comparator (AlphaFold2 [3], hereafter referred to by name) throughout this study. For every one of the 29 targets retained in the final comparison, this structure was an AlphaFold2 prediction: obtained directly from the AlphaFold Protein Structure Database where public coverage existed for the biologically relevant construct, or predicted in-house using AlphaFold2 and independently validated against the experimental reference prior to use (C*α* alignment RMSD *<* 0.5 Å, full sequence coverage) where public coverage was absent. No substitute structure predictor was used for any target in the final comparison.

### Proprietary Ensemble-Aware Pose Scoring

An in-house machine learning scoring architecture, integrated into the Mavchen-1 platform, was developed to evaluate protein–ligand poses generated across a conformational ensemble and to identify near-native binding poses without prior knowledge of the correct answer, drawing on the general class of graph-based neural network architectures established for structured molecular and biomolecular data [40–43]. The model architecture, training procedure, featurization scheme, and inference pipeline constitute proprietary technology and are not disclosed here. The model was trained and validated using a leave-proteins-out cross-validation scheme (10 folds, three held-out proteins per fold, with no held-out protein’s data present in any training fold) [44], ensuring that reported performance reflects generalization to unseen targets rather than memorization.

### Physical Validity and Accuracy Assessment

All final poses, from both the ensemble-derived and static-structure workflows, were assessed for physical and chemical plausibility using an independent, field-standard structural validity framework [19] encompassing bond geometry, steric clash, ring planarity, and protein–ligand contact/overlap criteria. Pose accuracy was quantified as symmetry-corrected heavy-atom RMSD [45, 46] to the experimentally determined native pose, computed in each structure’s own correctly aligned reference frame. A pose was classified as accurate at a conventional 2.0 Å RMSD threshold, consistent with established practice in the structure-based drug design literature [2].

### Statistical Analysis

Paired comparisons between the ensemble-based and static-structure approaches were conducted using the Wilcoxon signed-rank test [18] on per-target best-achieved RMSD. Success rates were defined as the fraction of targets meeting the RMSD accuracy threshold, evaluated both independently and under the combined requirement of RMSD accuracy and passage of all physical-validity checks. Category-stratified analyses were performed to determine whether performance differences were uniform across mechanistic classes or concentrated within specific categories, following cluster-stratified validation practice recommended for structure-based scoring benchmarks [44]. Model discrimination under class imbalance was additionally characterized using the area under the precision–recall curve, a metric more sensitive to minority-class performance than the area under the receiver operating characteristic curve alone [20]. All statistical tests were two-sided; significance was assessed at *α* = 0.05.

## Data and Code Availability

The protein target list, category assignments, and summary statistical outputs supporting the conclusions of this study are available upon reasonable request. The Mavchen-1 conformational sampling workflow, ensemble docking module, and proprietary pose-scoring model architecture, training data, trained model weights, and inference code are proprietary to Covenant Biosciences and are not made publicly available.

## Author Contributions

All authors contributed to study design, implementation, analysis, and manuscript preparation.

## Data Availability

The protein target list, category assignments, and summary statistical outputs supporting the conclusions of this study are available upon reasonable request to the corresponding author. The conformational sampling workflow, ensemble docking module, and proprietary pose-scoring model architecture, training data, trained model weights, and inference code are proprietary to Covenant Biosciences and are not made publicly available (see Methods, *Data and Code Availability*, for full detail).

## References

1. J. Lyu et al. Ultra-large library docking for discovering new chemotypes. Nature, vol. 566, pp. 224–229, 2019., 2019.

2. E. Lionta, G. Spyrou, D. K. Vassilatis, and Z. Cournia. Structure-based virtual screening for drug discovery: Principles, applications and recent advances. Curr. Top. Med. Chem., vol. 14, no. 16, pp. 1923–1938, 2014., 2014.

3. J. Jumper et al. Highly accurate protein structure prediction with AlphaFold. Nature, vol. 596, pp. 583–589, 2021., 2021.

4. R. Evans et al. Protein complex prediction with AlphaFold-Multimer. bioRxiv, 2021.10.04.463034, 2021., 2021.

5. Z. Lin et al. Evolutionary-scale prediction of atomic-level protein structure with a language model. Science, vol. 379, pp. 1123–1130, 2023., 2023.

6. M. Baek et al. Accurate prediction of protein structures and interactions using a three-track neural network. Science, vol. 373, no. 6557, pp. 871–876, 2021., 2021.

7. X. Cui, L. Ge, X. Chen, Z. Lv, S. Wang, X. Zhou, and G. Zhang. Beyond static structures: protein dynamic conformations modeling in the post-AlphaFold era. Brief. Bioinform., vol. 26, no. 4, p. bbaf340, 2025., 2025.

8. S. Vats, R. Bobrovs, P. Söderhjelm, and S. Bhakat. AlphaFold-SFA: Accelerated sampling of cryptic pocket opening, protein-ligand binding and allostery by AlphaFold, slow feature analysis and metadynamics. PLoS ONE, vol. 19, no. 8, p. e0307226, 2024., 2024.

9. P. Cimermancic et al. CryptoSite: Expanding the druggable proteome by characterization and prediction of cryptic binding sites. J. Mol. Biol., vol. 428, pp. 709–719, 2016., 2016.

10. A. Kuzmanic, G. R. Bowman, J. Juárez-Jiménez, J. Michel, and F. L. Gervasio. Investigating cryptic binding sites by molecular dynamics simulations. Acc. Chem. Res., vol. 53, no. 3, pp. 654–661, 2020., 2020.

11. D. E. Koshland Jr. Application of a theory of enzyme specificity to protein synthesis. Proc. Natl. Acad. Sci. USA, vol. 44, no. 2, pp. 98–104, 1958., 1958.

12. W. Sherman, T. Day, M. P. Jacobson, R. A. Friesner, and R. Farid. Novel procedure for modeling ligand/receptor induced fit effects. J. Med. Chem., vol. 49, no. 2, pp. 534–553, 2006., 2006.

13. P. Matricon, R. R. Suresh, Z.-G. Gao, N. Panel, K. A. Jacobson, and J. Carlsson. Ligand design by targeting a binding site water. Chem. Sci., vol. 12, no. 3, pp. 960–968, 2021., 2021.

14. J. Monod, J. Wyman, and J.-P. Changeux. On the nature of allosteric transitions: A plausible model. J. Mol. Biol., vol. 12, pp. 88–118, 1965., 1965.

15. J. H. Lin, A. L. Perryman, and J. A. McCammon. Computational drug design accommodating receptor flexibility: The relaxed complex scheme. J. Am. Chem. Soc., vol. 124, pp. 5632–5633, 2002., 2002.

16. R. E. Amaro, R. Baron, and J. A. McCammon. An improved relaxed complex scheme for receptor flexibility in computer-aided drug design. J. Comput. Aided Mol. Des., vol. 22, pp. 693–705, 2008., 2008.

17. R. E. Amaro, J. Baudry, J. Chodera, Ö. Demir, J. A. McCammon, Y. Miao, and J. C. Smith. Ensemble docking in drug discovery. Biophys. J., vol. 114, no. 10, pp. 2271–2278, 2018., 2018.

18. F. Wilcoxon. Individual comparisons by ranking methods. Biometrics Bull., vol. 1, no. 6, pp. 80–83, 1945., 1945.

19. M. Buttenschoen, G. M. Morris, and C. M. Deane. PoseBusters: AI-based docking methods fail to generate physically valid poses or generalise to novel sequences. Chem. Sci., vol. 15, no. 9, pp. 3130–3139, 2024., 2024.

20. J. Davis and M. Goadrich. The relationship between precision-recall and ROC curves. in Proc. 23rd Int. Conf. Machine Learning (ICML), 2006, pp. 233–240., 2006.

21. K. Zhu, C. Li, K. Y. Wu, C. Mohr, X. Li, and B. Lanman. Modeling receptor flexibility in the structure-based design of KRASG12C inhibitors. J. Comput. Aided Mol. Des., vol. 36, no. 8, pp. 591–604, 2022., 2022.

22. J. Abramson et al. Accurate structure prediction of biomolecular interactions with AlphaFold3. Nature, vol. 630, pp. 493–500, 2024., 2024.

23. Chai Discovery Team. Chai-1: Decoding the molecular interactions of life. bioRxiv, 2024.10.10.615955, 2024., 2024.

24. K. A. Kirou, M. Dall’Era, C. Aranow, and H.-J. Anders. Belimumab or anifrolumab for systemic lupus erythematosus? A risk-benefit assessment. Front. Immunol., vol. 13, p. 980079, 2022., 2022.

25. L. C. Arneson, K. J. Carroll, and E. M. Ruderman. Bruton’s tyrosine kinase inhibition for the treatment of rheumatoid arthritis. ImmunoTargets Ther., vol. 10, pp. 333–342, 2021., 2021.

26. Q. Ain, M. Batool, and S. Choi. TLR4-targeting therapeutics: Structural basis and computer-aided drug discovery approaches. Molecules, vol. 25, no. 3, p. 627, 2020., 2020.

27. H. M. Berman, J. Westbrook, Z. Feng, G. Gilliland, T. N. Bhat, H. Weissig, I. N. Shindyalov, and P. E. Bourne. The Protein Data Bank. Nucleic Acids Res., vol. 28, no. 1, pp. 235–242, 2000., 2000.

28. R. Wang, X. Fang, Y. Lu, and S. Wang. The PDBbind database: Collection of binding affinities for protein-ligand complexes with known three-dimensional structures. J. Med. Chem., vol. 47, no. 12, pp. 2977–2980, 2004., 2004.

29. P. Eastman et al. OpenMM 7: Rapid development of high performance algorithms for molecular dynamics. PLoS Comput. Biol., vol. 13, no. 7, p. e1005659, 2017., 2017.

30. J. A. Maier, C. Martinez, K. Kasavajhala, L. Wickstrom, K. E. Hauser, and C. Simmerling. ff14SB: Improving the accuracy of protein side chain and backbone parameters from ff99SB. J. Chem. Theory Comput., vol. 11, pp. 3696–3713, 2015., 2015.

31. J. Wang, R. M. Wolf, J. W. Caldwell, P. A. Kollman, and D. A. Case. Development and testing of a general amber force field. J. Comput. Chem., vol. 25, no. 9, pp. 1157–1174, 2004., 2004.

32. W. L. Jorgensen, J. Chandrasekhar, J. D. Madura, R. W. Impey, and M. L. Klein. Comparison of simple potential functions for simulating liquid water. J. Chem. Phys., vol. 79, pp. 926–935, 1983., 1983.

33. J. D. Chodera and F. Noé. Markov state models of biomolecular conformational dynamics. Curr. Opin. Struct. Biol., vol. 25, pp. 135–144, 2014., 2014.

34. O. Trott and A. J. Olson. AutoDock Vina: Improving the speed and accuracy of docking with a new scoring function, efficient optimization, and multithreading. J. Comput. Chem., vol. 31, no. 2, pp. 455–461, 2010., 2010.

35. J. Eberhardt, D. Santos-Martins, A. F. Tillack, and S. Forli. AutoDock Vina 1.2.0: New docking methods, expanded force field, and Python bindings. J. Chem. Inf. Model., vol. 61, no. 8, pp. 3891–3898, 2021., 2021.

36. G. Corso, H. Stärk, B. Jing, R. Barzilay, and T. Jaakkola. DiffDock: Diffusion steps, twists, and turns for molecular docking. in Proc. Int. Conf. Learning Representations (ICLR), 2023., 2023.

37. D. Santos-Martins et al. Meeko: Molecule parametrization and software interoperability for docking and beyond. J. Chem. Inf. Model., 2025, doi: 10.1021/acs.jcim.5c02271., 2025.

38. G. Landrum. RDKit: Open-source cheminformatics software. 2006. [Online]. Available: https://www.rdkit.org, 2006.

39. W. L. DeLano. The PyMOL molecular graphics system. Schrödinger, LLC, 2002., 2002.

40. M. Fey and J. E. Lenssen. Fast graph representation learning with PyTorch Geometric. in Proc. ICLR Workshop on Representation Learning on Graphs and Manifolds, 2019., 2019.

41. P. Veličković, G. Cucurull, A. Casanova, A. Romero, P. Liò, and Y. Bengio. Graph attention networks. in Proc. Int. Conf. Learning Representations (ICLR), 2018., 2018.

42. S. Brody, U. Alon, and E. Yahav. How attentive are graph attention networks? in Proc. Int. Conf. Learning Representations (ICLR), 2022., 2022.

43. C. Yang, E. A. Chen, and Y. Zhang. Protein-ligand docking in the machine-learning era. Molecules, vol. 27, no. 14, p. 4568, 2022., 2022.

44. C. Kramer and P. Gedeck. Leave-cluster-out cross-validation is appropriate for scoring functions derived from diverse protein data sets. J. Chem. Inf. Model., vol. 50, no. 11, pp. 1961–1969, 2010., 2010.

45. R. Meli and P. C. Biggin. spyrmsd: Symmetry-corrected RMSD calculations in Python. J. Cheminform., vol. 12, p. 49, 2020., 2020.

46. R. T. McGibbon et al. MDTraj: A modern open library for the analysis of molecular dynamics trajectories. Biophys. J., vol. 109, no. 8, pp. 1528–1532, 2015., 2015.

